# Identification of a First-In-Class Small Molecule CAPON Binder Using Affinity Selection-Mass Spectrometry Screening

**DOI:** 10.1101/2025.11.05.686689

**Authors:** Ashraf N. Abdo, Hossam Nada, Moustafa Gabr

## Abstract

NOS1AP (CAPON) is an adaptor protein of neuronal nitric oxide synthase (nNOS) correlated with Alzheimer’s disease progression, making it an attractive yet unexplored therapeutic target. To assess its chemical tractability, we employed affinity selection–mass spectrometry (AS-MS) to screen approximately 10,000 small molecules for CAPON binding, identifying 52 initial hits. These compounds were further evaluated for true binding interactions and potential autofluorescence or quenching effects using the Dianthus platform. Five compounds were selected for quantitative affinity determination by microscale thermophoresis (MST). Among these, compound **MA32** exhibited a dissociation constant (Kd) of 74 μM. **MA32** thus represents the first AS-MS–identified small-molecule binder of CAPON. This study establishes a workflow for small-molecule discovery against novel, previously uncharacterized targets.

## 1. Introduction

CAPON (NOS1AP) plays a role in regulating degenerative neurological diseases, neurotoxicity, heart diseases, and various other diseases. CAPON is an adaptor protein that binds to nNOS and regulates nitric oxide (NO) signaling in neurons[1-3]. CAPON is involved in modulating NMDA receptor-mediated signaling[4], and has roles in synaptic plasticity, apoptosis, and neuronal development [5]. CAPON has a C-terminal PDZ-binding motif, which can bind and interact with the N-terminal PDZ domain of nNOS. Thus, it is named the carboxy-terminal PDZ ligand of nNOS [6].

nNOS uses its PDZ domain to bind partner proteins, most notably PSD-95 and CAPON, which compete for the same binding site[7, 8]. Under physiological conditions, PSD-95 links nNOS to the NMDA receptor, forming a three-protein complex that activates nNOS after glutamate-induced calcium influx[2]. The resulting nitric oxide (NO) production supports normal neuronal signaling, synaptic plasticity, and learning. When glutamate levels become excessive, however, the same pathway drives overactivation of nNOS, generating toxic amounts of NO and peroxynitrite, which ultimately damages neurons[9, 10].

CAPON can interfere with displacing PSD-95 from nNOS, forming an alternative complex that dampens nNOS activity and reduces NO output, thereby offering neuroprotection against excitotoxic stress[11]. While this CAPON-mediated regulation is protective under pathological stimulation, CAPON overexpression can also inhibit normal PSD-95–nNOS signaling, which may negatively affect healthy neuronal function[9].

Neurodegenerative and psychiatric disorders such as Alzheimer’s[12], Parkinson’s disease, Huntington’s disease[13], amyotrophic lateral sclerosis[14], as well as conditions including major depressilion, bipolar disorder[15], anxiety, and PTSD [5], have been linked to dysregulation of the nNOS–CAPON interaction. When CAPON is overexpressed, it can trigger abnormal phosphorylation of Tau protein and promote formation of the Dexras1–nNOS–CAPON complex, a cascade that enhances β-amyloid accumulation and accelerates the progression of Alzheimer’s pathology[16, 17]. In more severe cases, this signaling imbalance contributes to synaptic failure and neuronal loss[12]. Alterations in CAPON-dependent pathways also affect downstream MAPK signaling, reducing ERK phosphorylation while activating p38, changes that ultimately drive neurotoxic outcomes [11, 18].

Further evidence comes from studies in the hippocampal dentate gyrus showing that disruption of the nNOS–CAPON complex influences the behavioral response to antidepressants such as fluoxetine. When CAPON remains bound to nNOS, levels of phosphorylated ERK, CREB, and BDNF are reduced, impairing neuroplasticity and increasing anxiety- and depression-like behaviors[19]. Because of these combined effects on molecular signaling and neuronal function, the CAPON–nNOS axis has emerged as a promising underexplored therapeutic target for both neurological and psychiatric diseases.

To determine whether CAPON can be targeted by small molecules, we utilized an affinity selection–mass spectrometry (AS-MS) workflow[20-23], a label-free technique that enables the direct detection of protein–ligand interactions in solution. This method is especially advantageous for proteins like CAPON, which lack convenient enzymatic readouts and are difficult to screen using traditional activity-based assays. From a chemically diverse library (Maybridge), the AS-MS screen yielded 121 preliminary hits that showed evidence of CAPON binding. These candidates were subsequently validated using complementary biophysical assays, including microscale thermophoresis (MST), to confirm interaction and measure binding affinity.

Through this multi-step validation pipeline, we identified a confirmed small-molecule binder of CAPON. Although the compound binds in the micromolar range, it represents the first ligand discovered for CAPON. The results establish that CAPON is chemically tractable and can be engaged by non-biologic agents, providing a starting point for structure-guided optimization. More broadly, this work illustrates the value of AS-MS as a discovery platform for challenging targets that lack enzymatic function and are otherwise challenging to interrogate with conventional screening technologies.

## 2. Methods

### 2.1 Affinity Selection Mass Spectrometry (AS-MS)

AS-MS Screening was performed as described in supplemental data. [21].

### 2.2. Single-dose Dianthus screening

Protein–ligand binding was initially evaluated by microscale thermophoresis (MST). Recombinant CAPON carrying a C-terminal His tag was fluorescently labeled using the RED-tris-NTA dye (NanoTemper Technologies) following the manufacturer’s protocol. Briefly, 200 nM protein was incubated with 100 nM dye in Phosphate Buffered Saline PBS supplemented with 0.05% Tween-20 buffer (pH 7.4) for 30 minutes at room temperature in the absence of light. Following labeling, the protein was diluted into PBST and combined with test compounds to yield final assay concentrations of 20 nM protein and 250 μM compound. Samples were incubated for 30 minutes at ambient temperature, briefly centrifuged, and subsequently analyzed on a Dianthus NT.23 Pico instrument. Assay buffer containing DMSO alone served as the negative control. Each measurement was performed in triplicate, and the reported values represent the averaged data.

To account for intrinsic compound fluorescence, parallel control reactions were prepared in which test compounds (250 μM) were diluted into the assay buffer without protein, incubated under identical conditions, and analyzed using the same readout settings. In addition, potential fluorescence quenching effects were assessed by incubating 10 nM RED-tris-NTA dye with each compound and recording the resulting signal.

### 2.3. Dose–response characterization using Microscale thermophoresis (MST)

Compounds that produced a clear binding response in the single-point screen were subsequently evaluated in a concentration-dependent MST assay. Twelve-point serial dilutions were prepared, spanning a starting concentration of 250 μM down to the low-nanomolar range, and mixed with labeled CAPON-His protein while maintaining a final DMSO concentration of 2%. After a 30-minute equilibration step at room temperature, samples were loaded into standard MST capillaries and measured on a Monolith NT.115 instrument, using medium-to-high infrared laser power and 60–80% LED excitation in the red detection channel. Dissociation constants (Kd) were calculated using the MO.Affinity Analysis package (NanoTemper Technologies).

### 2.4. Homology Modeling and Structure Preparation

The three-dimensional structure of the CAPON protein was predicted using homology modeling due to the absence of an experimentally determined structure. The SWISS-MODEL server was employed for model generation, utilizing the crystal structure of the GULP1 PTB domain-APP peptide complex (PDB ID: 6itu[24].1.A, chain A) as the template structure. The template exhibited 29.45% sequence identity with the target CAPON which falls within an acceptable range for homology modeling of structurally conserved domains.

The generated model was subjected to stereochemical quality assessment using Ramachandran plot analysis. The phi-psi dihedral angle distribution demonstrated that the majority of residues occupied the most favored regions (depicted in red), with additional residues in allowed regions (shown in yellow), indicating acceptable backbone geometry. The average model confidence score (QMEANDisCo) was 0.57 ± 0.07, suggesting moderate reliability of the predicted structure, particularly suitable for binding site identification and molecular docking studies. The most probable ligand-binding pocket on the CAPON homology model was identified using PrankWeb[25], a web-based tool for ligand-binding site prediction. The algorithm identified a prominent cavity on the protein surface (highlighted in red in Figure XC) characterized by physicochemical properties conducive to small molecule binding, including appropriate hydrophobic and electrostatic characteristics.

Molecular docking simulations were performed using the Schrodinger Maestro Suite 2024.4. to evaluate the binding modes and affinities of compound MA32 with the CAPON protein. The docking grid was centered on the top-ranked binding site predicted by PrankWeb[25]. Both compounds were docked into the predicted binding pocket, and the resulting protein-ligand complexes were analyzed for binding pose geometry, interaction patterns, and theoretical binding affinity. Protein-ligand interaction profiles were analyzed using the Discovery Studio Visualizer software. Two-dimensional interaction diagrams were generated to visualize hydrogen bonds, hydrophobic contacts, and other non-covalent interactions between the ligands and CAPON residues. Three-dimensional surface representations were rendered to illustrate the spatial occupancy of ligands within the binding pocket and the complementarity between ligand shape and pocket topology.

## 3. Results

### 3.1 Affinity Selection–Mass Spectrometry High Throughput Screening

To find small-molecule items that target CAPON, an affinity selection–mass spectrometry (AS-MS) approach was used, following our set ways. A screening of about 10,000 compounds from the ThermoScientific HitFinder collection was undertaken. This library is a varied chemical sample from the larger Maybridge Screening Collection, having been put together by grouping based on molecular fingerprints and a similarity measure to cover much of the drug-like chemical area. Each compound meets Lipinski’s rules for being drug-like and they are all at least 90% pure.

As was described before, the AS-MS process involved mixing the CAPON protein with groups of small molecules, which allowed for possible protein-ligand connections to form. Separation of these connections from the loose compounds was then achieved by using size-exclusion chromatography (SEC). The attached ligands were next let go, looked at with high-performance liquid chromatography (HPLC), and their identity found using time-of-flight mass spectrometry. From screening the 10,000-compound library,120 early hits were found, which is about a 1.2% hit rate. A manual review to remove reactive or unhelpful molecules resulted in the selection of 52 compounds for more testing.

To determine small-molecule ligands targeting CAPON, we used an affinity selection– mass spectrometry (AS-MS) strategy following our established procedures[21]. Approximately 10,000 compounds from the ThermoScientific HitFinder collection were screened. This library represents a chemically diverse subset of the broader Maybridge Screening Collection, curated through clustering based on Daylight molecular fingerprints and a Tanimoto similarity threshold of 0.7 to maximize coverage of drug-like chemical space. Each compound adheres to Lipinski’s criteria for drug-likeness (ClogP ≤ 5, no more than 10 hydrogen bond acceptors, 5 hydrogen bond donors, and molecular weight ≤ 500 Da) and exhibits a purity of at least 90%.

In the AS-MS procedure as previously described [21], CAPON protein was incubated with mixtures of small molecules to enable formation of potential protein–ligand complexes. These complexes were then separated from unbound compounds through size-exclusion chromatography (SEC). The bound ligands were subsequently released, analyzed by high-performance liquid chromatography (HPLC), and identified using time-of-flight mass spectrometry. Screening of the 10,000-compound library yielded 120 preliminary hits, corresponding to around 1.2 % hit rate. After manual curation to exclude reactive or poorly functionalized molecules, 52 candidates were selected for follow-up validation.

### 3.2 Primary screening by Dianthus

In order to further examine these hits for secondary validation assays. A single-concentration screen was conducted on the Dianthus platform employing Temperature-Related Intensity Change (TRIC) technology to detect direct protein–ligand interactions. In this assay, each compound was tested at a concentration of 200 nM against 10 nM his-tagged CAPON protein. A strict selection threshold was applied, defining active compounds as those showing signal responses exceeding three times the standard deviation of the negative control (protein plus dye only). Of the 52 compounds analyzed, 10 compounds demonstrated reproducible CAPON binding (Figure 2.A)

**Figure 1.**
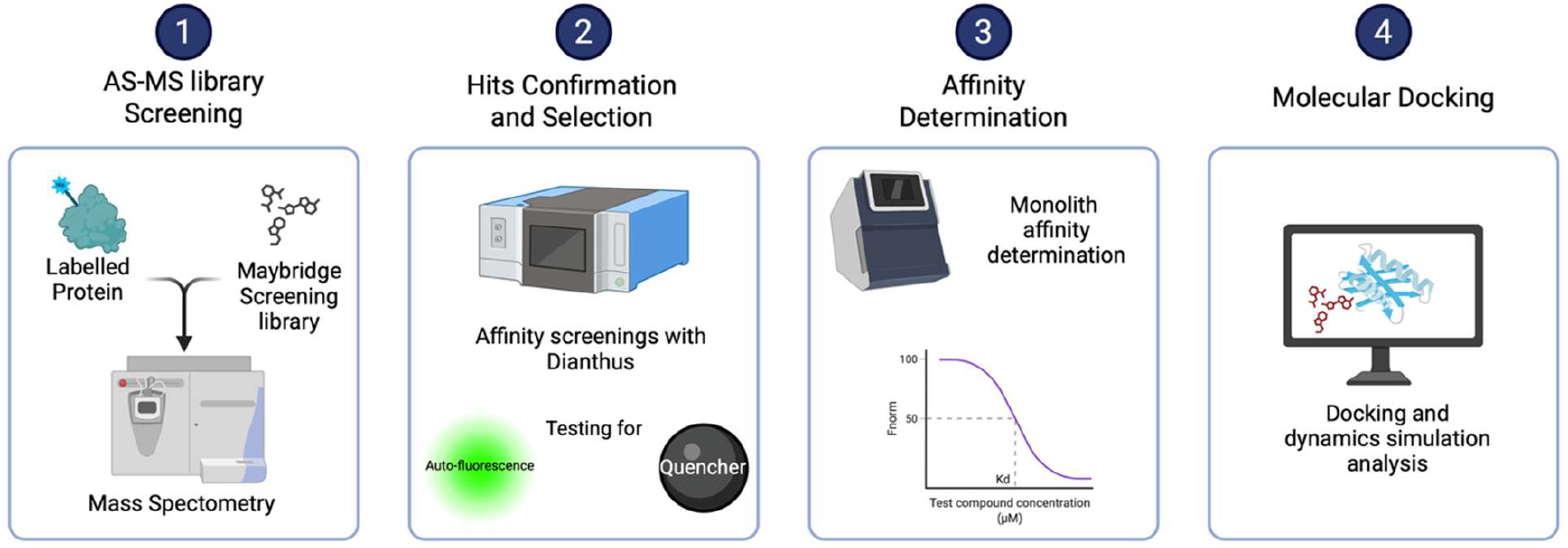
Schematic illustration of Experimental design for ASMS-based screening workflow for identifying small-molecule candidates as CAPON binders.

**Figure 2.**
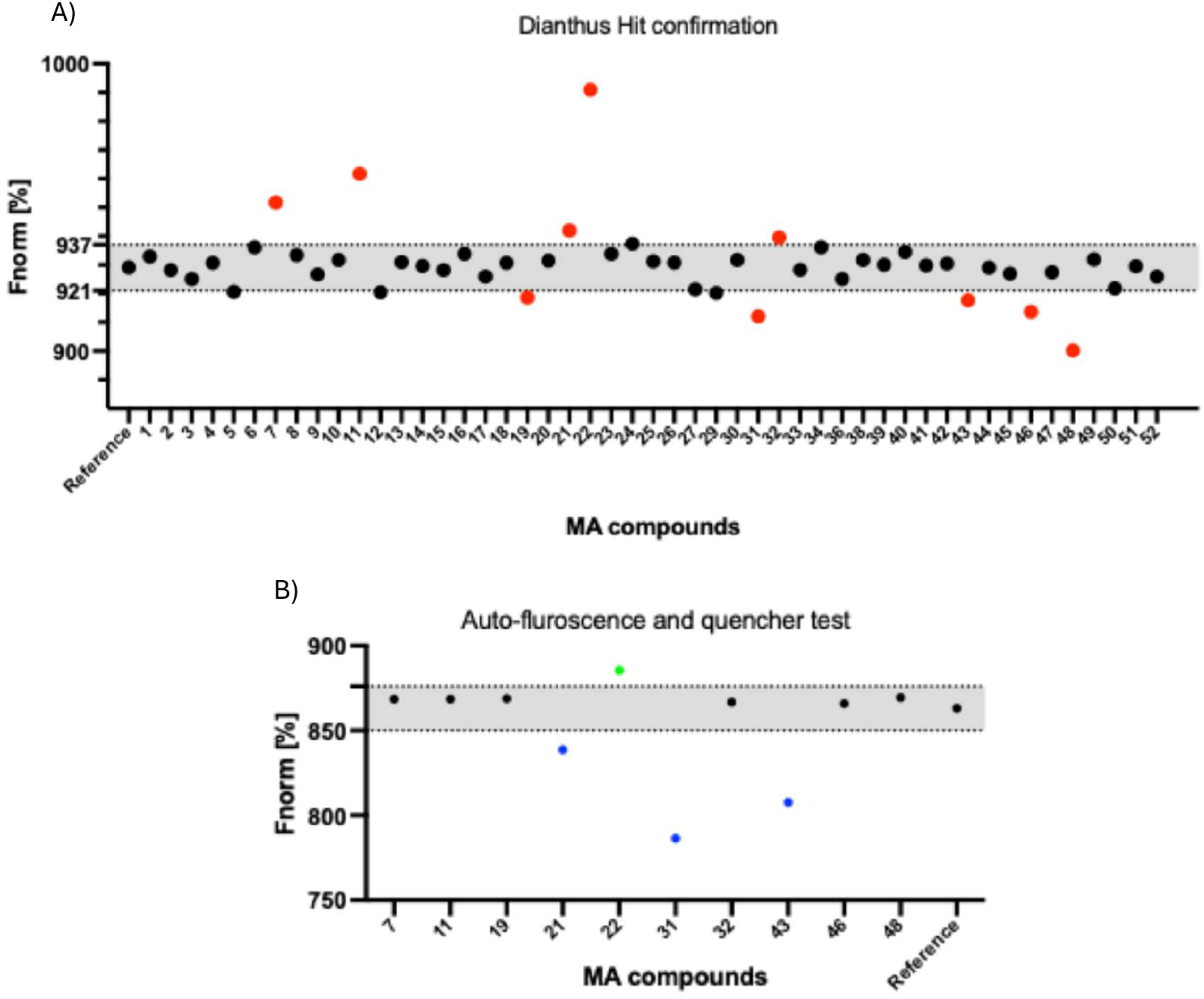
Dianthus Screening revealed potential hits. (A) Primary screening of compounds at 250 μM (with 2 % DMSO). Red dots indicate potential hits, negative control/reference (PBST buffer with 2 % DMSO). (B) Autofluorescence and quench test of candidates from single-dosage screening. Dotted lines in Fig. 2A and B indicate the threshold values (mean negative control (reference) ±3 ×SD) used to flag compounds with excessive autofluorescence

To check for potential signal interference, we compared the fluorescence of each compound to a reference sample containing only labeled CAPON and 2% DMSO. This analysis identified ten compounds (MA7, 11, 19, 21, 22, 31, 32, 43, 46 and 48) that showed fluorescence levels notably different from the reference (Figure 2.A). To eliminate the possibility of false positives caused by inherent compound fluorescence or interactions with the dye, we conducted additional control experiments, including autofluorescence quenching assay (Figure 2.B). This confirmatory test revealed that MA 22 displayed strong intrinsic fluorescence, whereas MA7, 11, 19, 32, 46 and 48 produced signals within the acceptable range in comparison to the reference samples. In the quenching assay, MA21, 31 and 43 showed a quenching effect and was consequently removed from further consideration. After this careful filtering process, only compounds that demonstrated minimal interference across all control tests were retained. Six candidates (MA7, 11, 19, 32, 46 and 48) met these criteria and were confirmed as reliable hits with consistent binding behavior.

### 3.3. Affinity determination by microscale thermophoresis (MST)

To verify binding interactions and determine affinity values quantitatively, compounds identified from the primary AS-MS screening were subjected to concentration-dependent binding analysis using microscale thermophoresis (MST). Fluorescently labeled His-tagged CAPON protein was titrated with serial dilutions of each candidate compound under optimized buffer (PBST) conditions containing 2% DMSO. The resulting thermophoretic responses were analyzed using the standard Hill model to calculate dissociation constants (Kd). Among the tested molecules, compound MA32 displayed a clear concentration-dependent shift, yielding a Kd of 74 ± 15 μM (Figure 3). This measurable binding affinity is consistent with fragment-like ligands typically identified through AS-MS–based discovery pipelines. Other compounds produced weak or inconsistent responses, suggesting low or nonspecific interactions. These results confirm that MA32 directly engages CAPON in solution, validating it as a bona fide small-molecule binder for further structural and functional evaluation.

**Figure 3.**
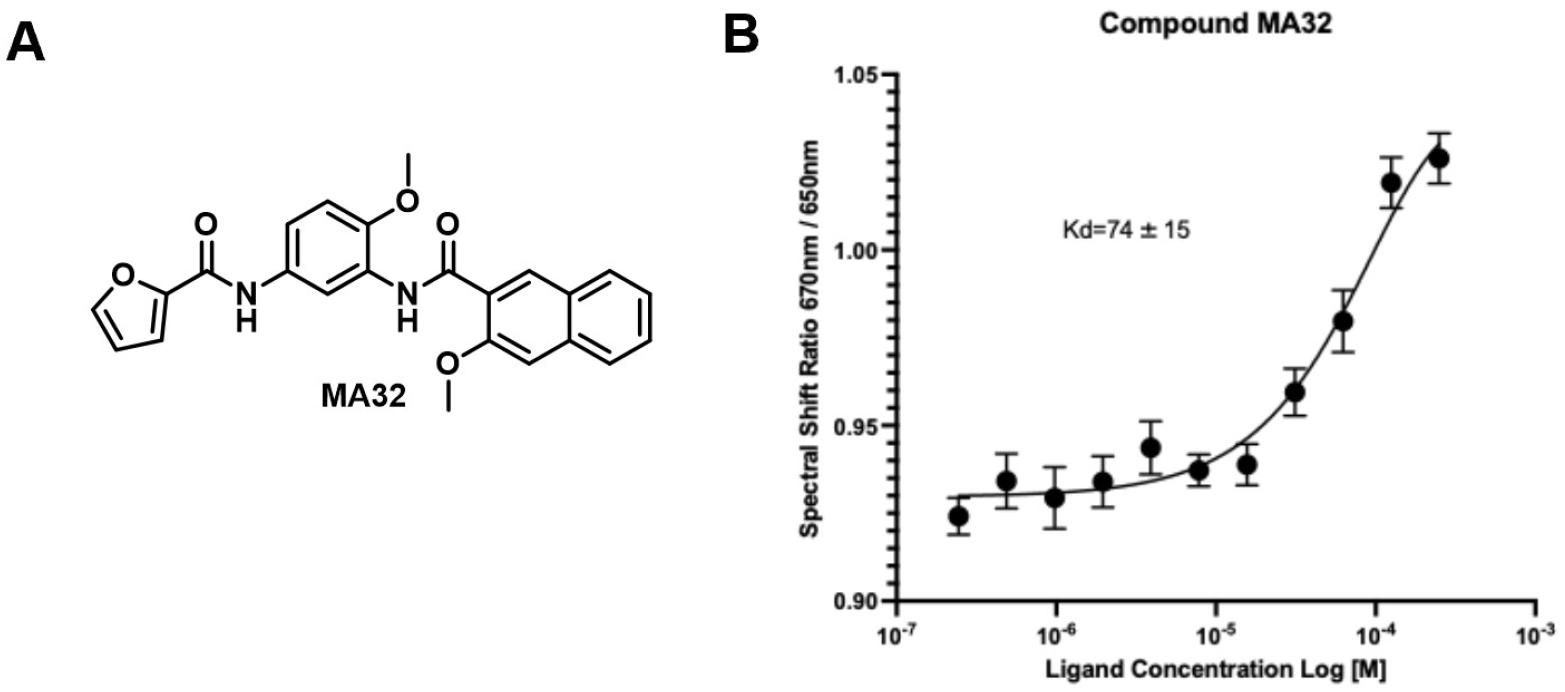
**A)** Chemical structure of MA32. **B)** Dose Response curve of Compound MA32 with CAPON protein using MST. The graph displays dose-dependent changes in spectral shift 670/650 nm ratio plotted against increasing concentration of the compound (n=3, Error bars display SEM).

### 3.4 Molecular Docking

The absence of an experimentally determined structure for CAPON necessitated the generation of a homology model using SWISS-MODEL, with the GULP1 PTB domain (PDB: 6itu.1.A) serving as the template despite modest sequence identity (29.45%). The resulting model exhibited acceptable stereochemical quality, with Ramachandran plot analysis demonstrating that the majority of residues occupied favored or allowed regions, and a QMEANDisCo score of 0.57 ± 0.07 indicating moderate structural reliability. PrankWeb analysis identified a druggable binding pocket characterized by a combination of charged, polar, and hydrophobic residues, which served as the target site for molecular docking studies. The successful docking of compound MA32 into this predicted binding site, with formation of multiple stabilizing interactions, suggests these molecules represent viable lead compounds for CAPON-targeted therapeutic development.

Detailed interaction analysis revealed that compound MA32 (Figure 4) engage the CAPON binding pocket through networks of hydrogen bonds, electrostatic interactions, and hydrophobic contacts, though with distinct binding signatures. MA32 demonstrates preferential engagement with LYS 65, LYS 69, TYR 36, LYS 70, VAL 108, and SER 39. The conserved interactions with LYS 65, 69, 70, TYR 36, and VAL 108 suggest these residues constitute a core binding motif essential for ligand recognition, while compound-specific interactions offer opportunities for structure-guided optimization to enhance potency and selectivity. The lysine-rich character of the binding site indicates that incorporation of acidic moieties or hydrogen bond acceptors may improve binding affinity in future derivatives.

**Figure 4.**
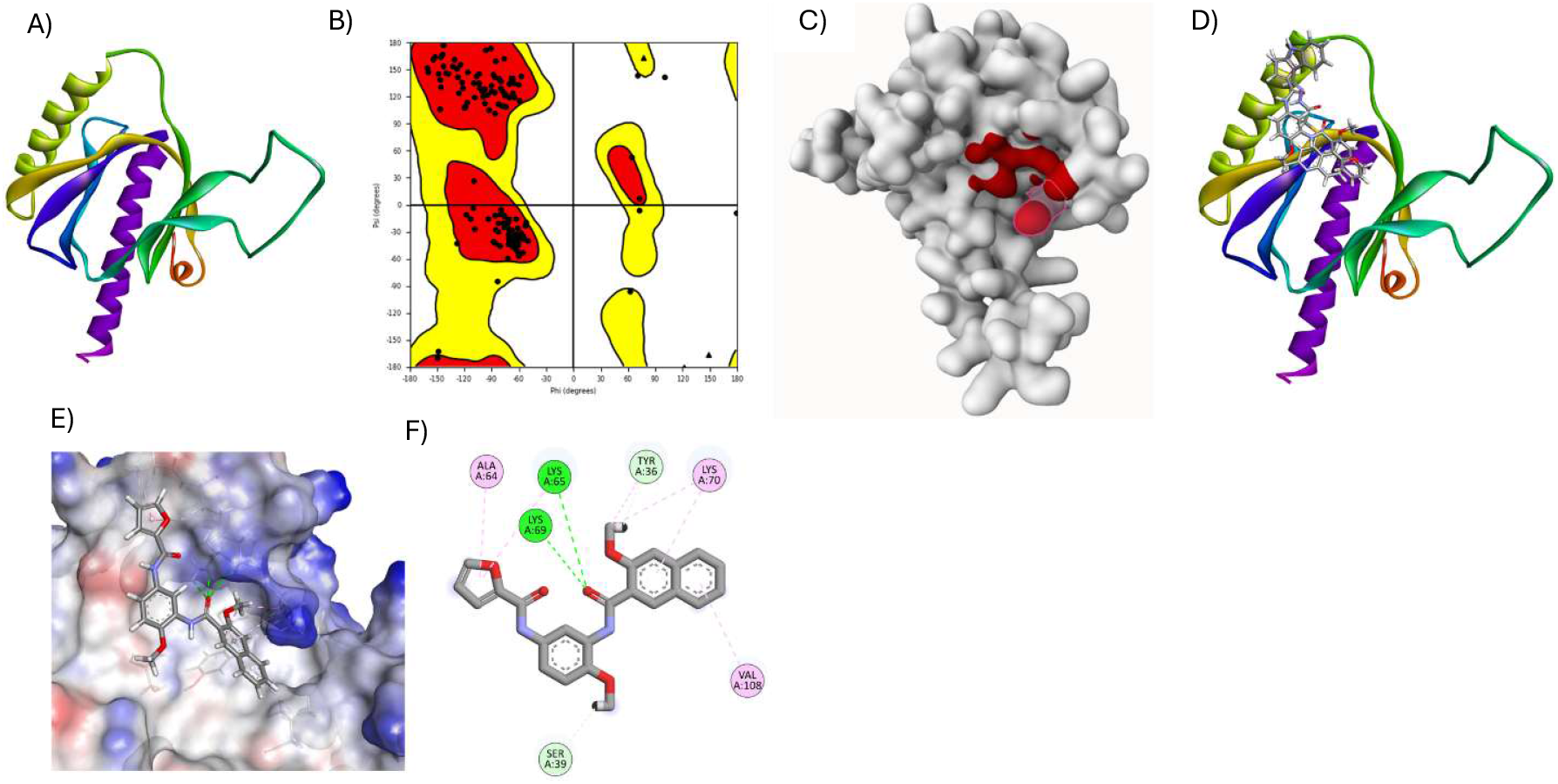
Homology modeling of CAPON and molecular docking analysis of compound MA32. (**A**) Cartoon representation of the CAPON homology model generated using SWISS-MODEL with the GULP1 PTB domain (PDB: 6itu.1. A) as template. (**B**) Ramachandran plot showing the distribution of phi-psi backbone dihedral angles for the CAPON model. Red regions indicate most favored conformations, yellow regions represent allowed conformations, and black dots denote individual residues. (**C**) Surface representation of the CAPON structure with the top-ranked ligand-binding pocket predicted by PrankWeb highlighted in red. (**D**) Overview of compound MA32 (shown as stick models) docked into the predicted CAPON binding site. (**E**) Detailed view of MA32 binding within the CAPON pocket. (**F**) Two-dimensional interaction diagram of MA32 bound to CAPON.

Several limitations constrain the interpretation of these computational findings. The moderate confidence score of the homology model introduces uncertainty regarding precise residue positioning and pocket geometry, which may affect the accuracy of predicted binding poses and interaction profiles. The reliance on static docking protocols does not account for protein flexibility, induced-fit effects, or the dynamic nature of protein-ligand recognition, which could substantially influence binding energetics and specificity. Furthermore, the predicted binding affinities lack experimental validation, and the functional consequences of MA32 binding to this site remain unknown. Future investigations should prioritize experimental validation through biophysical binding assays (surface plasmon resonance, isothermal titration calorimetry, or biolayer interferometry), determination of high-resolution co-crystal structures to confirm binding modes and functional studies to elucidate the biological impact of CAPON modulation by these compounds in relevant cellular contexts.

## Acknowledgements

This work was supported by the National Institutes on Aging under grant number RF1AG084635 (PI: Gabr). We would like to thank the Fisher Drug Discovery Resource Center of Rockefeller University (RRID: SCR_020985) for providing access to the Nanotemper Dianthus NT.23 Pico.

