## Supporting Information for "Identification of a First-In-Class Small Molecule CAPON Binder Using Affinity Selection-Mass Spectrometry Screening"

**Contents**

| **1.** | **Experimental for AS-MS screening** | **2** |
| --- | --- | --- |

**1. Experimental for AS-MS screening**

AS-MS Screening was performed using the Automated Ligand Identification System (ALIS) on an Agilent 2D-HPLC platform coupled to an Agilent time-of-flight (TOF) mass spectrometer. The assay was carried out using 0.1 mM recombinant human CAPON (His-tagged; SinoBiological) in PBS containing 300 mM NaCl and 0.01% Tween-20. The ALIS system integrates size-exclusion chromatography (SEC) driven by an Agilent 1260 HPLC pump with reversed-phase (RP) chromatography operated by an Agilent 1290 UHPLC pump, connected through a high-pressure switching valve to an Agilent 6230B TOF-MS. All spectra were collected in positive ion mode, and data were analyzed using Agilent MassHunter software in combination with proprietary computational tools.

SEC was performed using 700 mM ammonium acetate as buffer A and 70% acetonitrile as buffer B on a Polyhydroxyethyl A column (50 × 2.1 mm, 3 μm, 200 Å; PolyLC). RP chromatography was carried out using water with 0.1% formic acid (buffer A) and 90% acetonitrile with 0.1% formic acid (buffer B) on a Kinetex C18 column (50 × 2.1 mm, 2.6 μm, 100 Å; Phenomenex). For screening, a total of 10,000 compounds from the Maybridge HitCreator and HitFinder libraries were prepared in DMSO at 10 μM. Aliquots of 250 nL from each pooled mixture were dispensed into 384-well plates, diluted in assay buffer, and processed under ASMS conditions. A parallel LC-MS run was performed to verify that all compounds in the pools were detectable prior to affinity selection.

Data acquired from the screening campaign were processed using proprietary ASMS software, leading to the identification of 121 initial CAPON-binding hits. These results established the feasibility of the screening strategy and provided a prioritized set of candidate small molecules for follow-up biochemical and cellular validation studies.
